# SOAPy: a Python package to dissect spatial architecture, dynamics and communication

**DOI:** 10.1101/2023.12.21.572725

**Authors:** Heqi Wang, Jiarong Li, Siyu Jing, Ping Lin, Yu Li, Haibing Zhang, Yujie Chen, Zhen Wang, Hong Li

## Abstract

Advances in spatial omics technologies have brought opportunities to dissect tissue microenvironment, while also posing more requirements and challenges for computational methods. Here we developed a package SOAPy to systematically dissect spatial architecture, dynamics and communication from spatial omics data. Specifically, it provides analysis methods for multiple spatial-related tasks, including spatial domain, spatial expression tendency, spatiotemporal expression pattern, cellular co-localization, multi-cellular niches, and ligand-receptor-mediated and spatial-constrained cell communication. Applying SOAPy on different spatial omics technologies and diverse biological fields has demonstrated its power on elucidation of biological questions about tumors, embryonic development, and normal physiological structures. Overall, SOAPy is a universal tool for spatial omics analysis, providing a foundation for continued investigation of the microenvironment.

## Introduction

Spatially resolved transcriptomics has been crowned Method of the Year 2020 by Nature Methods^1^. Since then, more and more experimental methods for measuring expression levels of genes, proteins or metabolites in a spatial context have been developed. These technologies include barcode-based and imaging-based ones, which differ in resolution, accuracy and throughout^2,3^. The most widely used 10X Visium spatial transcriptomics measures thousands of genes in each 55μm spot that typically contains 1-10 cells^4^. And imaging-based methods reach more microscopic resolution, such as MIBI-TOF^5^ and PhenoCycler-Fusion^6^, both detecting dozens of proteins at subcellular resolution. Additionally, spatial multi-omics technologies that simultaneously measure multiple molecular types are emerging, e.g NanoString GeoMx DSP for 18000 RNAs and 140 proteins in the region of interest (usually >100 cells)^7^.

With the development of experimental methods, corresponding analysis pipelines have been designed for pre-processing raw data from specific experimental platforms, such as Space Ranger for 10X Visium and MCMICRO for multiplexed tissue imaging^8^. Methods adapted from single-cell RNA sequencing (scRNA-seq) data analysis could be used to perform standard dimensional reduction, clustering, cell type annotation and marker selection for spatial-omics data^9^ that do not require spatial information. And for low resolution spatial technologies, various deconvolution methods have been developed to impute cell-type composition from the mixture of cells.

After these pre-processing, downstream analyses are largely independent of experimental technologies, focus on the key feature of spatial omics: space. For example, identifying spatial variable genes^10–12^, detecting spatial domains^13^, inferring genes or cell-subtypes associated with spatial localization, and so on^3^. Earlier algorithms were often designed for one specific task, tools that fit in with various analysis tasks are becoming popular. A pioneer work Giotto not only builds a data pre-processing pipeline similar to scRNA-seq data analysis^14^, but also provides modules for spatial pattern detection, cell neighborhood analysis, and interactive visualization. Squidpy provides scalable analysis framework for both spatial neighborhood graph and image, along with an interactive visualization tool^15^. stlearn is another integrated package for spatial transcriptomic analysis, which adds the functions of spatial trajectories and pseudotime analysis^16^. Investigating the spatial organization of tissue microenvironment are important applications of spatial omics, which may gain new insights in various biological fields. However, the related analysis methods are scattered or lacking, a package for integrative analysis of microenvironmental spatial organization is in an urgent need.

To address this problem, we present a package SOAPy (Spatial Omics Analysis in Python) to jointly perform multiple tasks for dissecting spatial organization, including spatial domain, spatial expression tendency, spatiotemporal expression pattern, co-localization of paired cell types, multi-cellular niches, and cell-cell communication. SOAPy improves on previous tools in three main areas (**Table S1**): (1) Providing several alternative methods for most tasks to be suitable for complex and diverse biological tissues and various analysis requirements. (2) Offering a factor decomposition strategy for high-order spatial data to discover the major modes of variations in spatial, time, sample or others. (3) Proposing a new method to combine ligand-receptor expression and spatial locations to better infer short-range and long-range cell communications. We also applied SOAPy to a wide range of public datasets to demonstrate its general applicability and interpretability. SOAPy will be one of the fundamental packages for spatial omics analysis in Python.

## Results

### Overview of the SOAPy package

SOAPy is composed of four modules: ***Data Preprocessing, Molecular Spatial Dynamics*** containing *Spatial Tendency* and *Spatiotemporal Pattern* analysis, ***Cellular Spatial Architecture*** for analyzing *Spatial Proximity* and *Spatial Composition*, and ***Spatial Communication*** that combines spatial distance, expression level and interaction mechanism of ligand-receptors to infer cell interactions (**Figure 1**). In addition, SOAPy provides rich visualization capabilities for all of the analysis methods mentioned above.

**Figure 1.**
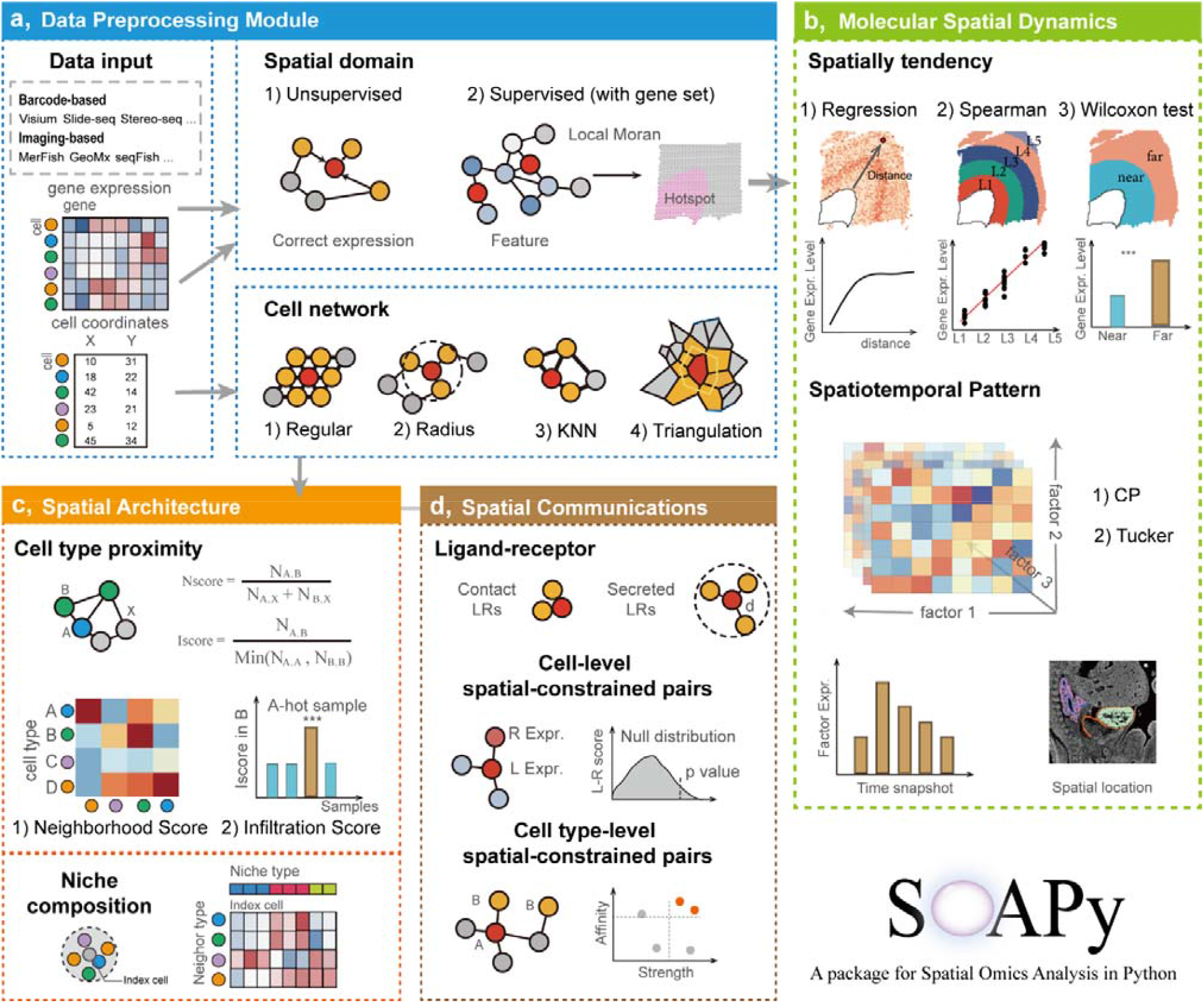
Schematic diagram of SOAPy. **a**, “Data Preprocessing” module that imports data, generates cell network and identifies spatial domains. Data from different spatial omics technologies are converted to a unified data structure. Cell network can be built by any of the four methods. Spatial domains are inferred by unsupervised learning from expression and morphological data, or supervised classification based on the expression of signature genes. **b**, “Molecular Spatial Dynamics” module. Spatial tendency analysis finds genes or cells whose expression change with spatial distance to the given region. **c**,Spatiotemporal Pattern analysis performs a tensor decomposition to discover the major modes of variation in space and time. **d**, “Cellular Spatial Architecture” module. Neighborhood and infiltration analysis find spatial proximal cell types. Spatial composition reveals conserved niches in which surrounding cells of the index cell are consisted of specific proportion of cell types. **e**, Innovative “Spatial Communication” module that combine spatial distance, expression level and action mechanism of ligand-receptors (LRs) to infer cell interactions. The contact and secreted LRs are considered for short-range and long-range cell communications, respectively. Results at cell/spot level indicate the heterogeneous interaction among different spatial locations, they are further integrated to cell type-level to report significant LRs for any two cell types.

The flexible ***Data Preprocessing*** module makes SOAPy suitable for various spatial data, fitting with different modalities and different resolutions. To demonstrate the utility of SOAPy, eight public datasets obtained from five state-of-the-art technologies were analyzed (**Table S2**). These datasets involve multiple scenarios with different molecular modalities (protein vs RNA), throughput (dozens to genome-wide), spatial resolution (0.1 ∼ 55μm), and in physiological and pathological states.

### Spatial domain analysis recapitulates anatomic and pathological structures

Cells are not randomly distributed in tissues. They are self-organized into specific structures to perform tissue functions. While in disease states, cells form abnormal structures. The *Spatial Domain* analysis provides unsupervised (STAGATE) and supervised (AUCell-LMI) methods to detect these structures (called spatial domains) based on gene expression profiles and spatial locations^13,17,18^.

We first tested STAGATE on Slide-seq V2 data for mouse olfactory bulb and 10x Visium spatial transcriptomic data for human breast cancer^19^. Spatial domains identified by STAGATE are highly consistent with the manual-labelled structures . It successfully distinguishes truth anatomical structures (**Figure S1a**), malignant and non-malignant tissues (**Figure S1b**, ARI=0.513), and more sophisticated pathological stages (**Figure S1c**, ARI=0.580). Then we tested AUCell-LMI for finding local structures with known signature genes, such as tertiary lymphoid structure (TLS)^20^. The results showed that supervised AUCell-LMI based on known TLS signature could more accurate and more convenient identified the TLS region than unsupervised STAGATE (**Figure S1d, e**). Taken together, Spatial domain analysis in SOAPy could extract the interesting anatomic or pathological structures for downstream analysis.

### Spatial tendency analysis finds genes associated with spatial structures

The aim of *Spatial Tendency* analysis is to assess whether expression features were influenced by spatial proximity to the region of interest (ROI). Expression features could be gene expression, pathway activity, cell proportion and so on. The ROI is defined by manual annotation or automatically detected by the *Spatial Domain* analysis. Two kinds of methods, statistical test and regression model, are available for tendency estimation in the *Spatial Tendency* module (Methods).

We used 10X Visium data of mouse dorsolateral prefrontal cortex (DLPFC)^21^ as an example to validate the feasibility of spatial tendency estimation (**Figure 2a**). The sample is consisted of the grey matter of DLPFC (including six cortical layers) and white matter (**Figure S2a**). To find genes whose expression changes along with the distance to the white matter, three strategies were used and compared^22^ (**Figure S2b, c**): 1) cortical layers were divided into two regions and applied Wilcoxon test to identify differential expressed genes; 2) cortical layers were separated to five continuous zones for Spearman correlation test; 3) a polynomial regression model was fitted between gene expressions and distances to the white matter. Some genes identified by Wilcoxon test and Spearman correlation only express in few spots, which may be the results of data sparsity instead of real biological differences (**Figure S2e**). The regression model describes the continuous spatial variation of expression, therefore it could find more complex spatial patterns than other methods^23^, such as nonlinear “low-high-low” spatial pattern (**Figure S2f)**. Next, we analyzed the expression patterns of 2857 significant (FDR < 0.05, range >0.3) genes identified by polynomial regression. K-means clustering grouped them into 10 clusters (**Figure 2b**). The gene clusters were compared with previously reported cortical layer specific genes^24,25^ (**Figure 2c**), showing high consistence. C3 is specifically highly expressed near white matter regions; the expression peaks of C5, C8, C2, and C7 are at layer 6, 5, 4, 2, respectively (**Figure 2d**).

**Figure 2.**
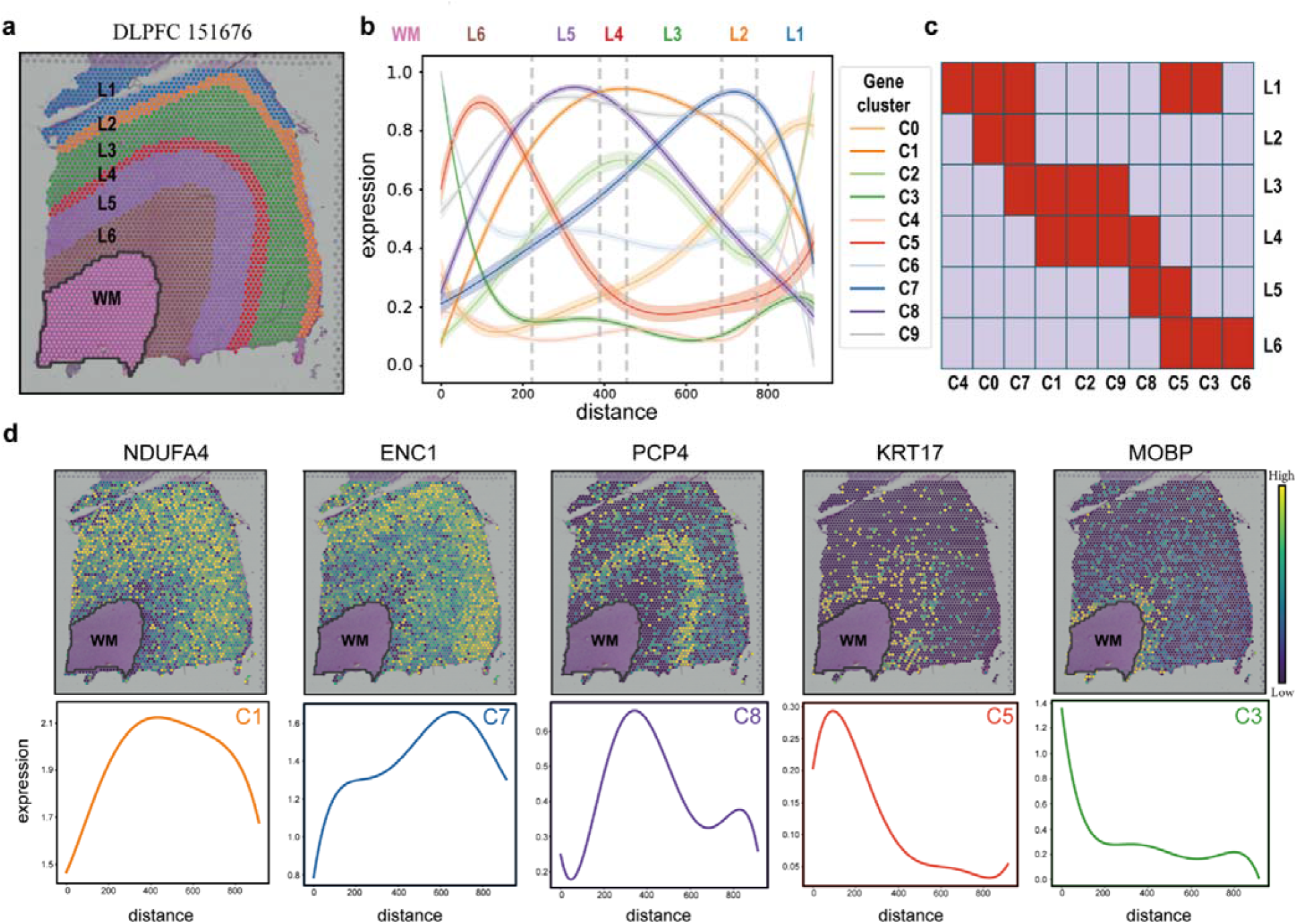
Spatial tendency analysis finds genes associated with spatial structures. **a**, HE image of a human dorsolateral prefrontal cortex (DLPFC) sample. Regions of white matter (WM) and six neuronal layers (L6 to L1) are labeled on the image. **b**, Regression curves between gene expression and the distance to WM. Polynomial regression models were fitted to identify genes whose expression varied along with the distance to WM boundary. These genes were grouped into 10 clusters by K-means clustering algorithm. Each curve present a cluster of genes with similar spatial expression tendency. Zero at the horizontal axis indicates the outer boundary of WM. **c**, Association between gene clusters and previously reported layer specific genes. Each row corresponds to a prior gene-list that specifically expresses in one neuronal layer^24^. Each red unit indicates the cluster of genes (column) is enriched in the prior gene-list (row). **d**, Spatial distributions and fitted curves of the representative genes.

Considering that there are no predetermined structures in some scenarios, we added three published methods (SpatialDE^10^, SPARK^12^, and SPARKX^11^) which identify spatial variable genes (SVGs) but do not need a given ROI. Comparing these SVGs methods with the above mentioned tendency estimation found shared and specific genes among methods (**Figure S2d**). SVG methods were more inclined to show the local differential expression of genes rather than the relationship with distance (**Figure S2g**). Users can select sutiable methods based on their requirements.

### Tensor decomposition reveals the spatiotemporal patterns of gene expression

With advances in omics techniques, spatial-resolved and time-series molecule profilings are becoming available. One of the challenges is how to study the roles of spatial effects and temporal effects simultaneously in biological questions. The *Spatiotemporal Pattern* function in SOAPy employs tensor decomposition to extract components from the three-order expression tensor (“Time-Space-Gene”), revealing hidden patterns and reducing the complexity of data explanation.

Here, we used the mouse embryo development dataset from GeoMx Digital Spatial Profiling (DSP)^7^. Limited by the availability of expression profiles, four time points (E9, E11, E13, E15) and eight subtissues (Heart wall, Heart valve, heart trabecula, Lung epithelium, Lung mesenchyme, Midgut epithelium, Midgut mesenchyme, and Midgut neuron) from three organs were included in our analysis (**Figure 3a,b**). Canonical Polyadic (CP) decomposition^26^ was used to factorize the expression tensor with 1000 high variable genes (a 4*8*1000 tensor) into seven factors, each of which is the outer product of three vectors that contain the loadings for describing the relative contribution of time, subtissues and genes (**Figure 3c**). We observed three empirical spatiotemporal patterns based on the loadings of time and subtissues: pure temporal variation (F1, F2), pure spatial variation (F3, F4), spatial and temporal variation occur together (F5, F6, F7). We also conducted functional enrichment analysis based on the loadings of genes for each factor (**Table S3**) and visualized the typical genes in images (**Figure 3d**).

**Figure 3.**
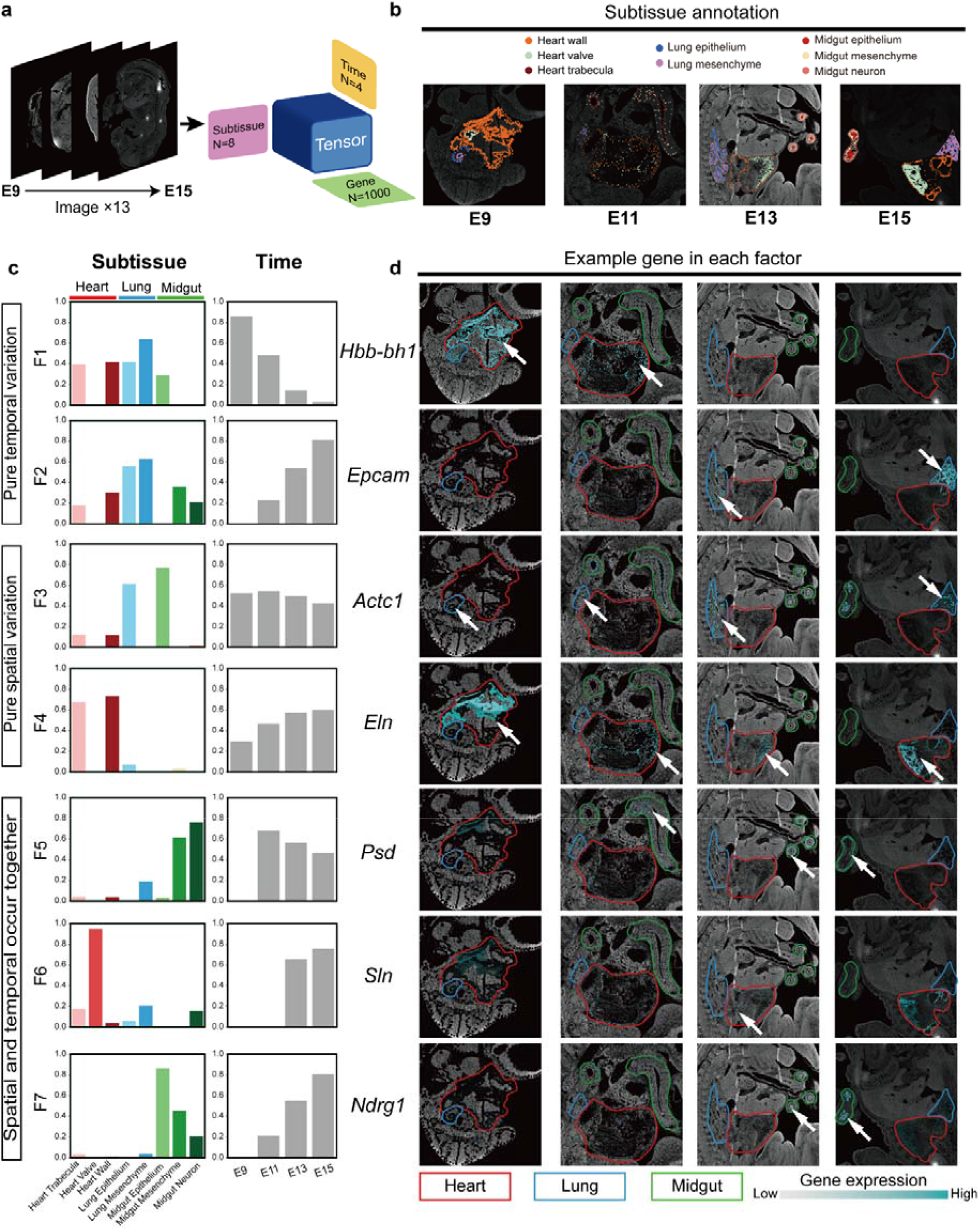
Tensor decomposition reveals the spatiotemporal patterns of gene expression during mouse embryo development. **a**, The spatiotemporal dataset of mouse development is represented by a three-order tensor (4 time points * 8 sub-tissues * 1000 highly variable genes), and then it’s decomposed into seven latent factors. **b**, Representative spatial locations of sub-tissues at four time points. Each spot in the subtissues represents an ROI. **c**, Loading vectors of space and time for each factor obtained by tensor decomposition. Higher loading values indicates larger contribution of sub-tissues or time points to the expression variation of this factor. **d**, Spatial expression of example genes for each factor. The contours of heart, lung and midgut are colored by red, blue and green curves. ROIs of gene expression are presented by cyan points. The darker the cyan color, the higher the gene expression level.

Genes in F1 (e.g. *Hbb-bh1*) highly express in heart and lung sub-tissues at E9, and then gradually decrease in the later stages. Their expression pattern is consistent with the enriched function “regulation of vasculature development”. F1 indicates co-development of heart and lung in the early embryo, which is consistent with previous studies^27^. The expression of F2 genes (e.g. *Epcam*) increases significantly since E11 in most sub-tissues of three organs, especially in the lungs. Expression of F3 and F4 genes is stable along the developmental time. F4 genes highly express in heart wall and heart trabecula, and their functions are enriched in cardiac cell development as expected. Both F5 and F7 genes are enriched in midgut development. F5 (e.g. *Psd*) slightly decreases from E11 to E15, while F7 (e.g. *Ndrg1*) increases obviously from E11 to E15. F6 genes are specifically highly expressed in the heart valve between E13-E15. In summary, the *Spatiotemporal Pattern* function in SOAPy could reveal spatiotemporal specificity during development and other biological processes.

### Spatial proximity analysis characterizes co-localization patterns between cell types

Spatial architecture of cells is important for understanding the organization rules from single cells to tissues^28–30^. SOAPy first constructs a cell/spot network fromspatial locations; then implements two scenarios for deciphering spatial architecture: *Spatial Proximity* analysis (including neighborhood and infiltration) determines whethe two cell types or cell states within an image are significant proximal; *Spatial Composition* analysis identifies multi-cellular niches that composed by cell types with specific proportion.

We applied this analysis to a dataset of 41 triple-negative breast cancer (TNBC) patients^5^, which used multiplexed ion beam imaging by time-of-flight (MIBI-TOF) to simultaneously quantify expression of 36 proteins in-situ at sub-cellular resolution. Totally 211,649 cells were annotated to eight types (epithelial cell, endothelial cell, mesenchymal cell, B, CD4 T, CD8 T, macrophage and other) based on the expression of known protein markers.

First, *Spatial Neighborhood* analysis was performed to identify significantly adjacent cell types compared to random perturbation^29^. **Figure 4a** illustrates the neighborhood score of all samples for all cell type pairs, with positive or negative scores corresponding to co-localization or avoidance. Different immune cells types such as B, CD4 T, CD8 T and macrophage have significant co-localization in many patients, which may relate with the formation of inflammatory foci (**Figure 4b**). Endothelial and mesenchymal cells also prefer to co-locate together (**Figure 4c**). Colocalization pattern of malignant epithelial cells and non-parenchymal cells were highly heterogeneous across patients. Taking malignant epithelial cells and mesenchymal cells as an example, samples with less than 200 mesenchymal cells were filtered, others are subjected to *Spatial Infiltration* analysis. Samples with higher and lower infiltration scores indicate mixed (e.g. sample 28) and compartmentalized (e.g. sample 29) patterns between malignant epithelial cells and mesenchymal cells respectively (**Figure 4 d-f**).

**Figure 4.**
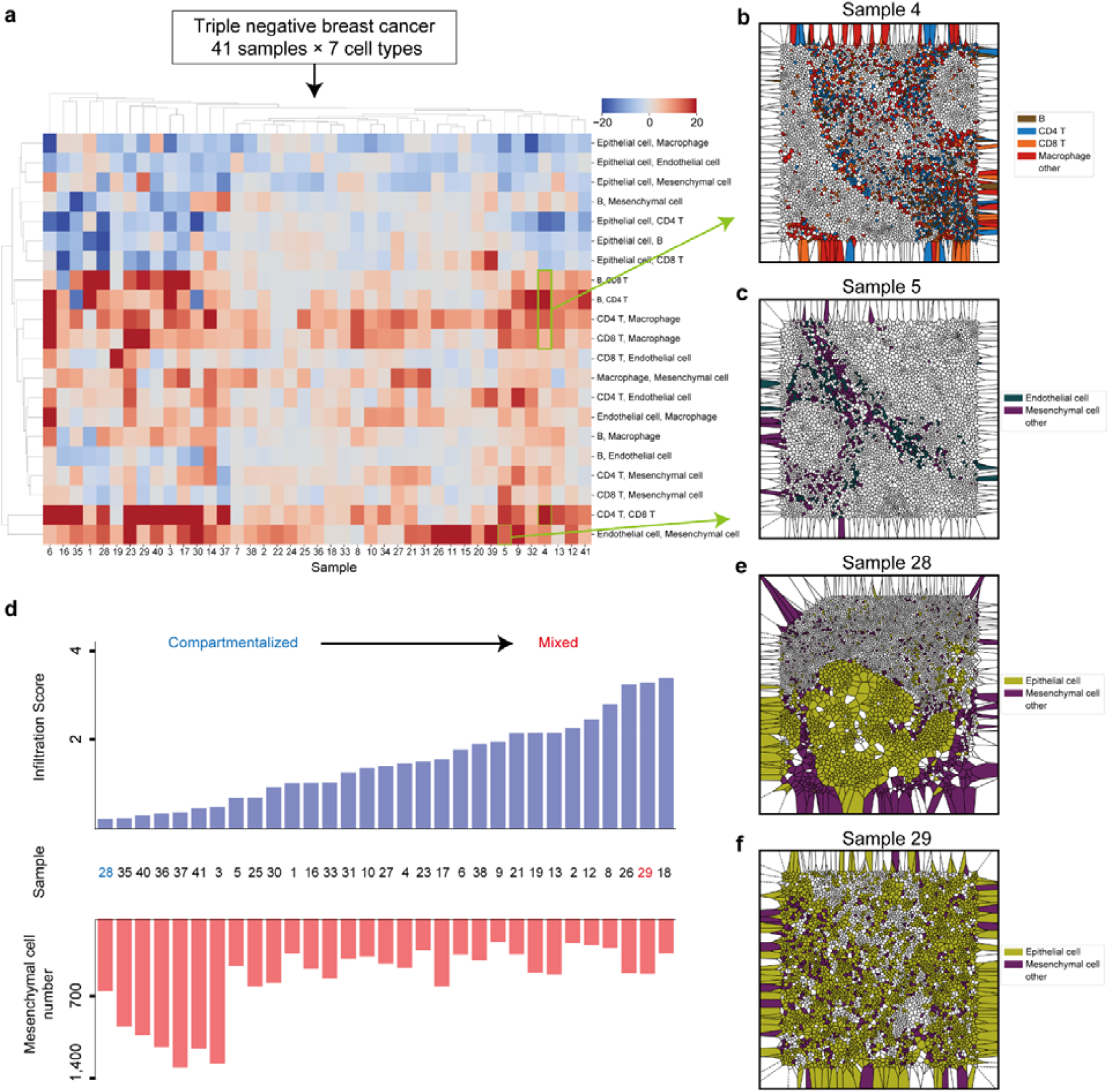
Spatial proximity analysis characterizes cellular co-localization patterns. The triple negative breast cancer (TNBC) dataset contains 41 samples and 7 cell types. **a**, Heatmap showing the neighborhood scores of any two cell types in all TNBC samples. **b**, A representative sample with strong co-localization among immune cells. **c**. A representative sample with strong co-localization between endothelial and mesenchymal cells. **d**, The red bars show the number of mesenchymal cells and the blue bars show the infiltration score of mesenchymal cells into malignant epithelial cells. **e**, A representative sample with low infiltration score, suggesting compartmentalization between mesenchymal cells and tumor tissues. **f**, A representative sample with high infiltration score, suggesting mixture of mesenchymal cells into malignant epithelial cells.

### Spatial composition analysis discovers multi-cellular niches

For *Spatial Composition* analysis of the TNBC dataset, the cell-cell network that connected centroids of the cells within 100 pixels was built to capture the composition pattern of more surrounding cells. Niche of each cell was presented by the proportion of cell types of its surrounding cells, called I-niche. I-niches of 211,649 cells from 41 TNBC patients were clustered into 30 niche clusters, named C-niches (**Figure 5a, Figure S3a**). The major cell types of the top two C-niches (C-niche13, C-niche18) are mainly composed of malignant epithelial cells, and the percentages of other cell types are less than 15%, showing the characteristics of tumor cell aggregation (**Figure 5b**). Additionally, epithelial cells also form C-niches with other cell types. For example, C-niche25 is composed of 38% epithelial cells, 31% mesenchymal cells, and 9% macrophages; C-niche27 is composed of 23% epithelial, 28% endothelial, 10% mesenchymal cells and 10% macrophages; C-niche15 is composed of 30% epithelial, 23% CD4 T, 13% CD8 T cells and 11% macrophages, suggesting different local microenvironment exists among tumors (**Figure 5b**). We also observed four B cell dominated C-niches (C-niche10, C-niche17, C-niche28, C-niche4) that may be related to tertiary lymphoid structures. For example, sample 1 contains C-niche 10, 17, and 28 (**Figure 5c**). Around 80% of cells are B cells in C-niche10; C-niche17 majorly consists of 52% B cells, 13% CD8 T cells, 10% CD4 T cells, and 11% epithelial cells; C-niche28 majorly consists of 30% B cells, 10% CD8 T cells, and 37% epithelial cells.

**Figure 5.**
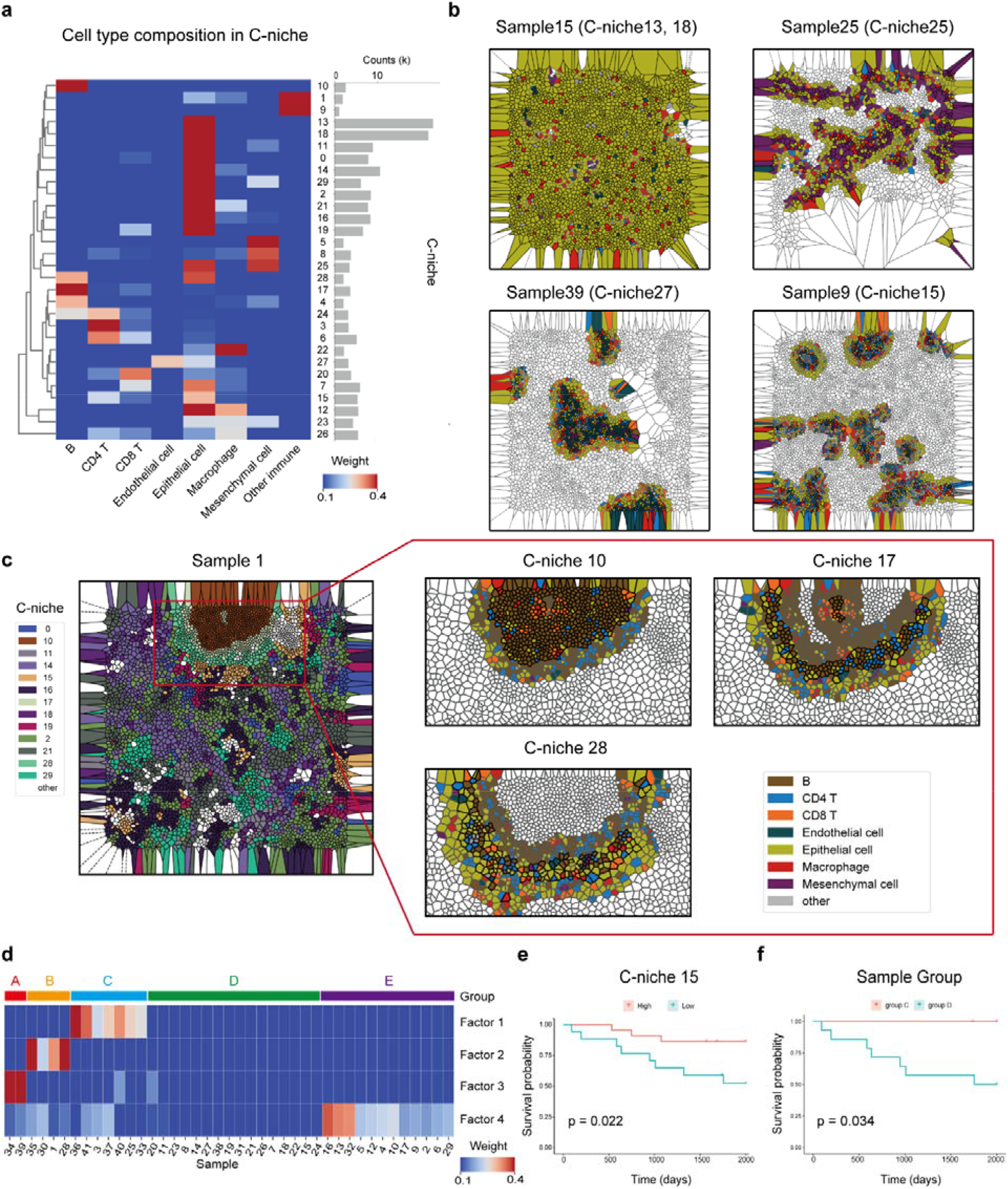
Spatial composition analysis discovers multi-cellular niches in TNBC samples. **a**. Heatmap on the left shows the composition of neighbor cells in each C-niche. The right barplot shows the number of cells belonging to each C-niche. **b**, Representative samples of different C-niches, characterizing tumor cell aggregation and different local microenvironment of tumors. **c**, The left image shows an example sample that has B cell dominated C-niches (the region of red box). Cells are colored by C-niches. ‘other’ are low-frequent c-niches whose proportion is less than 2%. Right images are amplified views of three representative C-niches. Black or gray cell contours indicate cells belonging to or not belonging to the C niche. The fill colors of cells represent cell types involved in the definition of the C-niche. **d**, Heatmap showing the loading values and clusters of samples. The three-order ‘Niche-CellType-Sample’ tensor was decomposed to four latent factors (**Figure S3b, c**). Samples are clustered into five groups according to their loading vectors. **e**, Survival curves stratified by the proportion of C-niche-15. **f**, Comparison of survival curves between two groups of patients.

In order to investigate the combinational effects of non-parenchymal cell types and niches on patient heterogeneity, the “Niche-CellType-Sample” tensor (30*7*41) was factorized to four factors (**Methods**). All samples were clustered into five groups according to the sample loadings in different factors (**Figure 5d**). Sample groups A, B, C, and E have the highest loadings in factors 3, 2, 1, and 4, respectively. By checking the loadings of cell types and niches in the major factors (**Figure S3b**,**c**), group B corresponds to the above mentioned B cell enriched samples; group C is characterized by niches with high proportion of mesenchymal cells; group E has niches consisted of T cells and macrophages.

Furthermore, survival analysis was performed to explore the clinical indications of niches. Eight c-niches were significantly related to survival time (P < 0.05, **Figure S4**). For example, patients with a higher proportion of c-niche15 had a longer survival time (**Figure 5e**). There also exists survival differences among the patient groups identified by the “Niche-CellType-Sample” tensor decomposition, such as longer survival time for group C patients that that of group D (**Figure 5f**). Taken together, spatial composition analysis could find multi-cellular niches and yield insight into how cells are organized into tissues.

### Ligand-receptor-mediated and spatial-constrained cell-cell communications

The above spatial architecture analysis disregards interacting molecules and context, while expression-based methods like CellphoneDB^31^ and CellChat^32^ infer cell-cell communications by the expression of ligands and receptors (LRs) disregarding spatial proximity. SOAPy develops a new method that simultaneously utilizes spatial location and gene expression to calculate interaction scores (affinity and strength) and then outputs significant LR interactions (**Figure 6a, Methods)**. It can not only infers short-range cell communication that relies on contact LRs to directly deliver signaling between adjacent cells; but also infer long-range cell communication that does not require cell–cell contact, rather depending on the diffusion of signaling molecules from one cell to another after secretion^33,34^.

**Figure 6.**
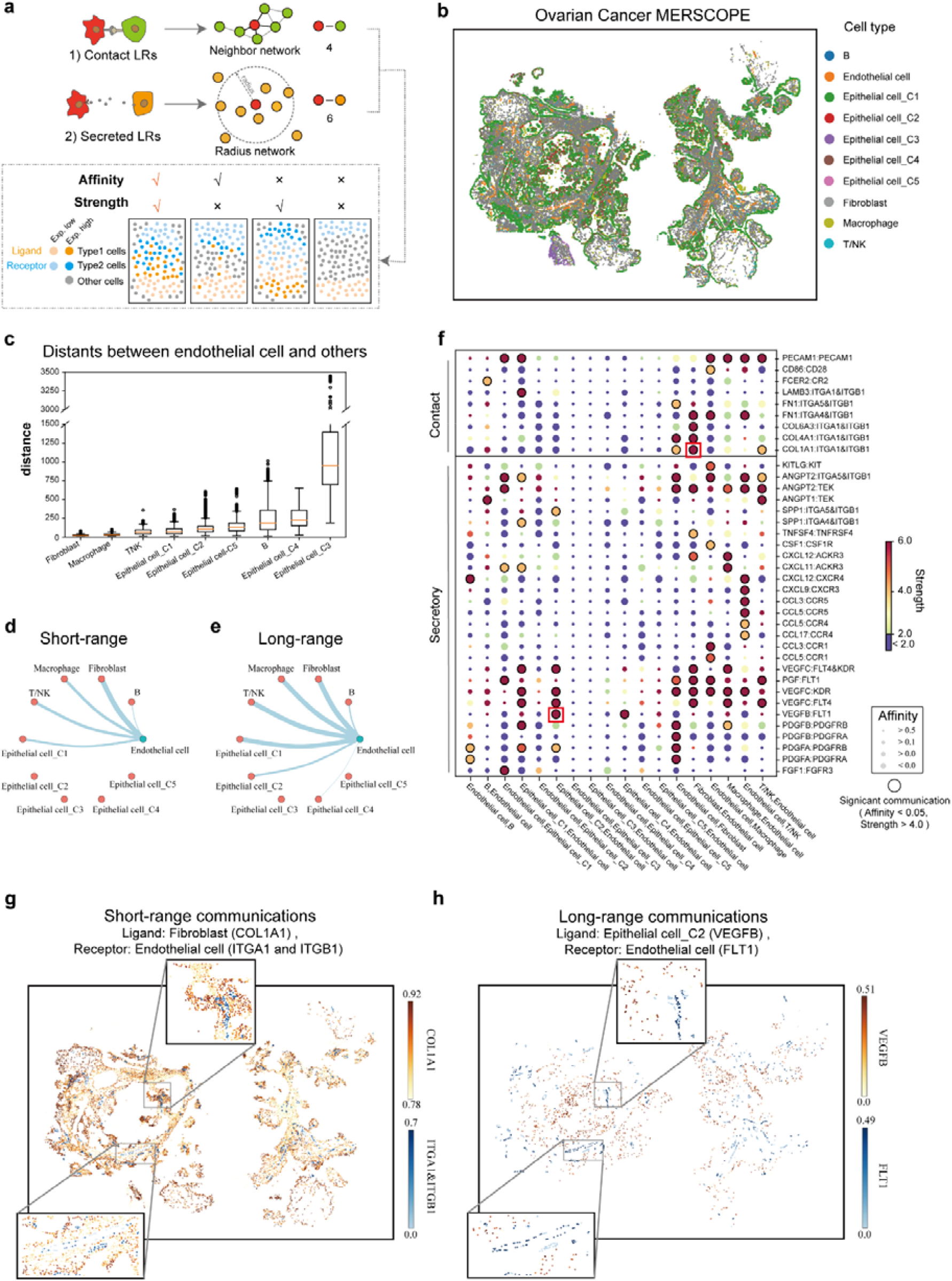
Ligand-receptor-mediated and spatial-constrained cell-cell communications. **a**, The brief flow chart of our method. Short-range interaction is mediated by contact LRs on neighbor cells, long-range interaction is mediated by secreted LRs on cells within the given radius. Two new metrics, affinity and strength, are defined to estimate the probability of LR interactions in any two cell types. Only when both metrics are high, the LR is significant to mediate the interactions of these two cell types. **b**, MERSCOPE data from an ovarian cancer sample. **c**, Barplot showing the shortest distance from other cell to the closest endothelial cell. **d, e**. Short-range and long-range cell communication networks between endothelial cells and other cell types. Edges in d and e are the number of contact and secreted LRs. Edge width is the number of significant ligand-receptor pairs (affinity P-value < 0.05, strength > 4). **f**, Dot plot with ligand-receptor interactions corresponding to d and e. Each row indicates a ligand-receptor pair, with the first and the second genes representing a ligand and a receptor, respectively. Dot size indicates P-value of affinity. Color indicates the strength score. **g**, An example of contact LR that mediates the communication between spatially colocalized fibroblast and endothelial cells. COL1A1 is the ligand on sender fibroblast cells, and ITGA1/ITGB1 is the receptor on receiver endothelial cells. Expression was scaled to the range of 0-1 by normalization. **h**, An example of secreted LR, corresponding to the communication between spatially separate epithelial and endothelial cells. VEGFB is the ligand on sender epithelial-hypoxia cells, and FLT1 is the receptor on receiver endothelial cells.

The *Spatial Communication* module was applied to an ovarian cancer dataset generated by the MERSCOPE platform, measuring 500 genes and 71,381 cells (**Figure 6b**). Cells were classified and annotated into ten types or subtypes by Leiden clustering algorithm. The spatial locations of epithelial cells C3 are very special, which clearly separated with most of other cells. Therefore, our method did not find significant contact LRs between epithelial cells C3 and other cell types. However, CellChat, one of the most popular LR communication inference packages using scRNA-Seq data, reported many LR interactions due to lack of spatial constrain (**Tabls S4**), indicating lower false positives of our method.

We used endothelial cell as an example to present its short-range and long-range communication partners. Fibroblasts and macrophages are located closest to endothelial cell, while epithelial cell C3 and C4 are far away from endothelial cell (**Figure 6c**). Consistently, fibroblasts have the largest number of contact LRs with endothelial cells recognized by our algorithm, while there is no contact LRs for distant cell types such as epithelial cells C2, C3, C4 and C5 (**Figure 6d**). For cell types that are not spatially close to endothelial cells, *Spatial Communication* module could infer secreted LRs that mediate long-range cell communications. The average distance from epithelial cells C2 to the closet endothelial cells is significantly larger than the average distance from fibroblasts to the closet endothelial cells (P < 3.9e-312). There are no contact LRs between epithelial cells C2 and endothelial cells but 6 secreted LRs were identified (**Figure 6 d, e**).

Totally, we found 19 contact LRs and 66 secreted LRs that may play key roles in short-range and long-range communication between endothelial cells and others (**Figure 6f**). For example, COL1A1 (type I collagen) and its receptor ITGA1/ITGB1 (integrin α/β) highly express on spatial adjacent fibroblasts and endothelial cells, their affinity and strength scores are significantly higher than random scores (**Figure 6g**). Previous studies have reported that binding collagen to integrin may activate downstream signaling pathways contributing to cancer progression^35^. VEGFB-FLT1 is an interesting LR pair for long-range communication between epithelial and endothelial cells (**Figure 6h**). Epithelial cells C2 release ligand VEGFB, and endothelial cells high express FLT1 (also known as VEGFR1). Their interaction may promote tumor angiogenesis and are potential drug targets for anticancer therapy^36^. In summary, SOAPy provide a new way to study spatial-constrained cell-cell interactions and more accurately identify the related ligand-receptor pairs.

## Discussion

Tissue microenvironment is critical for understanding homeostasis, development, regeneration and disease. Single-cell and spatial resolved omics are the most promising technologies to investigate microenvironment. Tools for systematically dissecting microenvironment and discover biologically important genes or spatial cellular architecture are still falling behind, SOAPy just fill this gap. SOAPy contains easy-to-use analysis modules for interpreting complex spatial microenvironments, such as the spatial distribution patterns of genes and cells, dynamic changes along with space and time, and cell-cell communications et al. In this article, we demonstrated all SOAPy modules with various types of spatial omics data, and provides complete tutorials to help users get started quickly.

The spatial distribution of genes or cells is associated with many elements, such as time, interaction of cells, pathological foci, sample heterogeneity and so on. In the face of these multi-dimensional data, how to extract important and meaningful features is a key task. SOAPy utilizes tensor decomposition to discover the major modes of variations from multi-dimensional data. The cases of mouse embryo development and breast cancer showed that tensor decomposition in SOAPy is powerful for interpret complex biological data. Another significant advantage of SOAPy is the innovative *Spatial Communication* module. It combines spatial distance, expression level and interaction mechanism of ligand-receptors to infer cell-cell communication. The case of ovarian cancer showed that SOAPy could markedly reduce false positives of interacting ligand-receptors compared to existing methods.

These advantages makes SOAPy differ from existing spatial data analysis tools. Future extensions of SOAPy could be the integration of multi-modal spatial data to delineate microenvironment, adaptation of methods from geoscience, network science, or artificial intelligence to better extract biological meaningful spatial patterns. We anticipate that SOAPy will be widely used by researchers to discover biological insights from spatial omics data.

## Methods

### Data preprocessing

#### Data Import

The *Data Import* function converts data from different spatial omics technology to a unified data structure that contains expression profiles of molecules (genes/proteins/metabolites) and location of cells/spots. Barcode-based data formats can be read directly by passing in tables representing expression matrix and spatial coordinate information. An image and a cell segmentation mask are provided for imaging-based data, and the representation and coordinate matrix is extracted through the tutorials on our website. We used the Scanpy toolkit^37^ and generate Anndata data.

#### Spatial network construction

The *Spatial network* function provides four ways to build a neighborhood network of cells/spots (Figure 1a). 1) Regular network; 2) KNN network that connects each site with its K nearest neighbors; 3) Radius network that all cells/spots within the given distance are connected; 4) Neighbor network based on Voronoi Diagram.

### Spatial domain identification

#### Unsupervised spatial domain identification: STAGATE

STAGATE is a graph attention autoencoder for spatial domain identification^13^. It firstly integrates gene expression profiles and spatial location information to learn low-dimensional latent embedding, and then assigns spatial domains by Louvain clustering.

#### Supervised spatial domain identification: AUCell-LMI

To detect domains whose signature genes are already known, the score of signature genes for each cell/spot is calculated by AUCell^38,39^, and then local Moran index^17^ (LMI) is used to estimate the degree of spatial aggregation. LMI of cell/spot *i* is defined as:

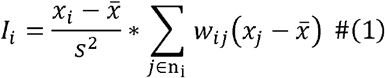

Where *x*_*i*_ is the AUCell score of cell/spot, 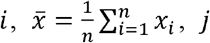, is any neighbor cells/spots of *i* based on K nearest neighbors,*w*_*ij*_ is the spatial weight between and *j*. The P-value is calculated by permutation test and adjusted by Benjamini-Hochberg method^40^ to get the false discovery rate (FDR).

LMI of all cells/spots are illustrated by Moran scatterplot (Figure S1e). Each point represents one cell/spot, the horizontal axis shows the normalized AUCell score, and the vertical axis indicates the “spatial lag” which is calculated by spatial weighted normalized score of neighboring sites. Sites with positive AUCell scores, positive spatial lags, and low FDR were picked out as the targeted spatial domain.

### Spatial tendency analysis

#### Definition of ROI and distance

Given a region of interest (ROI), the first step is to generate a binary mask file (**Figure S2a**). Users can manually select ROI using tools like ImageJ to generate a mask file, or get interesting cells/spots via SOAPy *Spatial domain* analysis and then use SOAPy to create a mask file: Discrete cells/spots are converted to continuously connected regions using a series of digital image processing steps in OpenCV library, such as dilation, corrosion, removal of small connected components, and removal of holes.

Next, the shortest distance from each cell/spot to the ROI boundary (contour) is calculated. When an ROI contains multiple connected components, the closest connected component is selected to calculate the distance^23^.

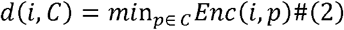

where *i* is a cell/spot, *C* is the boundary of ROI, and *p* is any pixel on the boundary. *Enc*( )is a function of Euclidean distance. Distance with positive or negative signs are used respectively to distinguish cells/spots located outside or inside the ROI boundary. Then we can study the tendency of molecule expression along with distance.

#### Identification of expression features with spatial tendency

SOAPy provides two statistical testing methods (**Figure S2b**): 1) wilcoxon rank sum test to compare the molecule expression of cells/spots between two regions; 2) spearman correlation between median expression and the rank of continuous zones. To resolve more complex spatial tendency (e.g., nonlinear) or analyze ROIs without prior hypothesis, SOAPy provides a parameter regression method (polynomial regression model) and a non-parametric regression method (locally weighted liner regression, LOESS).

Polynomial regression assumes that the output variable can be represented by the sum of powers of the input variable.

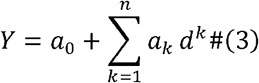

Where *d* is the distance to the ROI; Y is the vector of molecule expression; *n* is the degree of the polynomial; *a*_0_ is intercept; *a*_*k*_ are slope coefficients. P-value is calculated by F-test.

LOESS is a locally weighted polynomial regression method. Its core concept is to fit weighted linear regression models with each data point using its surrounding data points within the predefined window size and connect the centers of the regression lines. *R*^2^ (coefficient of determination) and residual standard deviation are estimated to measure the goodness of fit.

Parameters used in both of the regression models could be customized and adjusted based on the biological scenario and goodness of fit. To summarize the spatial tendency of all molecules, the estimated expression values are fed into the K-means clustering algorithm to obtain gene clusters with similar spatial expression tendency.

### Spatial architecture analysis

#### Spatial neighborhood analysis

For each paired cell types, a neighborhood score (*NS*) between cell type 1 (*ct*1) and cell type 2 (*ct*2) is calculated as follows^29^:

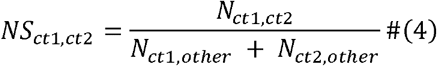

where *N*_*ct*1,*ct*2_ is the number of direct connections between *ct*1 and *ct*2, *N*_*ct*1,*other*_ is the number of direct connections between *ct*1 and all other cell types. Background distribution is generated from 1000 random permutations that fix the numbers of *ct*1 and *ct*2 and randomly change their locations. P-value is the proportion of permutations whose *NS* is larger or smaller than the observed one, which corresponds to either avoidance or interaction between *ct*1 and *ct*2.

#### Spatial infiltration analysis

An infiltration score (*IS*) is defined to present the degree of non-parenchymal (immune or stromal) cells infiltration into malignant tissues:

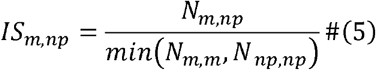

where *N*_*m,np*_ is the number of direct connections between malignant cells and non-parenchymal cells. Sample with too few non-parenchymal cells are regarded as cold tumor. Otherwise, larger infiltration score indicates more non-parenchymal cells are mixed into malignant tissues, while smaller infiltration score suggests non-parenchymal cells are more possible to be compartmentalized with malignant tissues.

#### Spatial composition analysis

Given an index cell, niche is defined as the proportion of cell types for its surrounding cells^41^. Taken all cells in one or more images, clustering algorithms like K-means divides their niches into different clusters, called C-niches.

#### Spatial-constrained cell-cell communication inference

Ligand-receptor (LR) pairs were obtained from the CellChat^32^ package, in which LR pairs were classified into contact and secreted based on their action mechanism. We hypothesized that the contact LR pairs mediate short-range cell communications while secreted LR pairs could mediate long-range cell communications. Therefore, SOAPy infers cell communications based on the types of LR pairs and spatial distance among cells (presented by a cell network). For short-range communication, direct neighbors on Voronoi Diagram are connected to build a cell network. For long-range communication, all cells within the given distance are connected to build a cell network. Once the cell network is built, *Affinity* and *Strength scores* scores are calculated for LRs on any two cell types. The LR pairs with *Affinity pvalue* < 0.05 and *Strength* > 4.0 are considered to be significant. Paired cell types are ranked based on the number of significant LRs.

#### Cell-level ligand-receptor affinity score

The interaction of LR is variable among cells/spots at different spatial locations, therefore we first define a cell-level ligand-receptor affinity score. Suppose a cell/spot *i* is a sender of ligand, cells/spots that have connection with *i* and express the matched receptor are receivers, the *Affinity scores* of ligand-receptor at location *i* is defined as:

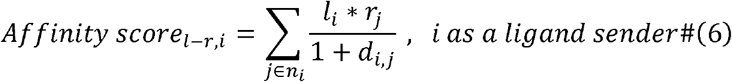

where *j* is the cell/spot that connect to *i* in the cell network; *l* and *r* are expression levels of the ligand and receptor; *d* is 0 for contact LR pairs or Euclidean distance between *i* and *j* for secreted LR pairs. Similarly, when the cell/spot *i* is a receptor receiver, the *Affinity scores* of receptor-ligand at location is defined as:

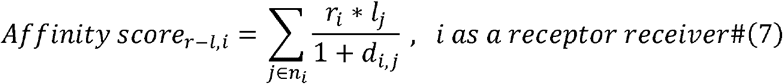

The *Affinity pvalue* is obtained by random permutation:

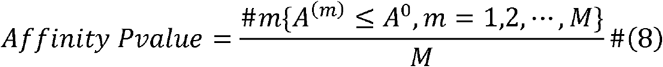

*M* is the total number of randomizations, *A*^(*m*)^ is the *Affinity scores* under the *m*-th randomization. Each randomization redistributes the expression values of the LR, but keeps topology of the cell network. The affinity scores are calculated for all cells/spots, and the P-values are used to find a subset of cells/spots at which the LR exist interaction.

#### CellType-level communication score

Suppose *ct*1 and *ct*2 are cell types that express ligands and receptors, respectively. The *Affinity scores* between the ligand of *ct*1and the receptor of *ct*2 is the sum of cell-level scores:

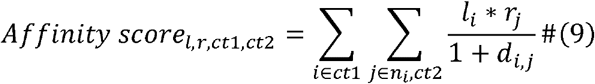

*Affinity pvalue* is also calculated by random permutation, which randomly assign a pseudo expression value to each cell/spot based on cell-type specific expression distribution.

*Affinity* reflects whether spatial connected *ct*1 and *ct*2 relatively more highly express the LR genes. However, if the expression of ligand or receptor is too low in *ct*1 or *ct*2 compared to other cell types, it is difficult to say that the LR is important for cell communications; Additionally, If *ct*1 and *ct*2 are connected by too few edges in the cell network, their communication may be false positive even affinity is significant. To address these problems, another index ‘strength’ is added. *Strength*_*l,r, ct*1,*ct*2_ consists of two components: one is the relative expression level of LR pairs on *ct*1 and *ct*2, and the other indicates the enrichment of real spatial connections between *ct*1 and *ct*2. The detailed definition is as follows:

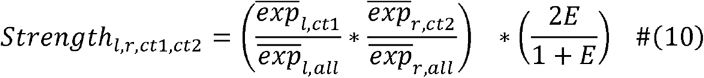

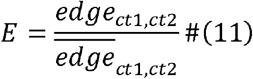

where 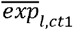 and 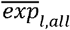 are the average expression of ligand in *ct*1and in all cells; *edge*_*ct*1,*ct*2_ and 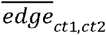 are the real and expected number of connections between *ct*1 and *ct*2; *E* is the ratio of real and expected numbers. To constrain the range of *E* and make the result more stable, a Hill function transforms *E* into a range of (0, 2) and keeps the transformed is *E* still 1 when the number of real and expected connections are equal.

### Tensor decomposition

To discover the major modes of variation in the high-order spatial data, such as the “Time-Space-Gene” tensor or “Niche-CellType-Sample” tensor, SOAPy provides interface functions to conveniently build tensors from AnnData objects and then decomposes tensors into several latent factors or components.

SOAPy implements two tensor decomposition methods, CANDECOMP /PARAFAC (CP) and Tucker decomposition^26,42^. Moreover, SOAPy supports non-negative constraints to make the factors more interpretable. Take non-negative CP^43^ as an example, an n-order tensor X is expressed as the weighted sum of R (user-defined number of factors) rank-one tensors:

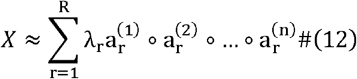

where λ is the weight of each factor; 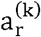 is the non-negative loading values of k-th variable in the r-th factor, indicating the relative contribution of variables to factors. Each factor is the outer product of the loading vectors.

### Availability

All data and code that produced the findings of the study, including all main and supplemental figures, are available at https://github.com/LiHongCSBLab/SOAPy.

## Acknowledgements

We acknowledge Andrew E. Teschendorff (from Shanghai Institute of Nutrition and Health, Chinese Academy of Sciences) for his advice on our manuscript. We thank Bihan Shen, Xi Yan (from Shanghai Institute of Nutrition and Health, Chinese Academy of Sciences) and Biao Liu (from Center for Excellence in Molecular Cell Sciences, Chinese Academy of Sciences), for their help on programing and result interpretation.

## Supplementary Information

**Figure S1.** Spatial domain analysis recapitulates anatomic and pathological structures. **a**, Anatomical structure of mouse olfactory bulb (Slide-seq V2 data) and domains identified by STAGATE. **b-c**, Expert-annotated pathological regions of a breast cancer sample (10x Visium), and the estimated 2-class and 19-class domains based on the results of by STAGATE. **d**, Expert-annotated tertiary lymphoid structure (TLS) on a kidney cancer sample (10x Visium), and the estimated TLS by the AUCell-LMI method. **e**, Moran scatterplot. The x-axis is the Z-transformed AUC, which presents the activity for the signature genes of TLS. The y-axis is the spatial weighted normalized AUC scores of neighboring locations. Hotspot presented by red points (FDR < 0.05, x > 0, y > 0) is regarded as tertiary lymphoid structure.

**Figure S2.** Spatial tendency analysis. **a**, Steps of image per-processing to generate a binary mask file for the given region of interest (ROI). **b**, Illustration of three spatial tendency analysis strategies: wilcoxon test, spearman correlation, and regression. **c**, Venn diagram shows the overlap of top 1000 genes (FDR q-value < 0.05) obtained from three spatial tendency analysis strategies. There are 380 overlapped genes, 352, 209 and 227 genes uniquely identified by a method (**Figure S2c**). **d**, Intersection plot showing the agreement for seven methods. Four methods estimate the tendency of gene expressions changing with the distance to a given region: wilcoxon test, spearman correlation, polynomial regression and LOESS regression. Other three methods identify spatially variable genes (SVGs) whose expressions depend on their spatial locations: SPARK, SPARKX and SpatialDE. The top-ranked genes with equal number obtained from each method were compared. Genes of LOESS are ranked by R-square, and genes of the remaining methods are ranked by FDR values. **e-g**, Representative genes identified by different kinds of methods. **e**: MIAT that is significant by Wilcoxon test and Spearman correlation analysis but not significant by regression methods; **f**: PVALB that is significant by regression methods; **g**: Expression of TFF1 is spatially variable but do not show tendency of change with the distance to WM.

**Figure S3. a**, Heatmap showing the proportion of niches in all TNBC samples. **b-c**, Loadings of cell types and niches obtained from the decomposition of “Niche-CellType-Sample” tensor.

**Figure S4.** Results of survival analysis.

**Table S1.** Comparison with existing tools of spatial omics data analysis.

**Table S2.** Examples datasets that were used in this study.

**Table S3.** Enriched functional terms by gene set enrichment analysis. Genes were pre-ranked based on the loading values of each factor obtained from tensor (“Time-Space-Gene”) decompositon.

**Table S4**. Predicted LR interactions between spatial-separated epithelial cells C3 and other cell types by CellChat and SOAPy.

